# The Fisher-Wright model with deterministic seed bank and selection

**DOI:** 10.1101/035246

**Authors:** Bendix Koopmann, Johannes Müeller, Aurélien Tellier, Daniel Živković

**Affiliations:** Center for Mathematics, Technische Universität München, 85748 Garching, Germany; Center for Mathematics, Technische Universität München, 85748 Garching, Germany and Institute for Computational Biology, Helmholtz Center Munich, 85764 Neuherberg, Germany; Section of Population Genetics, Center of Life and, Food Sciences Weihenstephan, Technische Universität München, 85354 Freising, Germany

## Abstract

Seed banks are a common characteristics to many plant species, which allow storage of genetic diversity in the soil as dormant seeds for various periods of time. We investigate an above-ground population following a Fisher-Wright model with selection coupled with a deterministic seed bank assuming the length of the seed bank is kept constant and the number of seeds is large. To assess the combined impact of seed banks and selection on genetic diversity, we derive a general diffusion model. We compute the equilibrium solution of the site-frequency spectrum and derive the times to fixation of an allele with and without selection. Finally, it is demonstrated that seed banks enhance the effect of selection onto the site-frequency spectrum while slowing down the time until the mutation-selection equilibrium is reached.

## INTRODUCTION

Dormancy of reproductive structures, that is seeds or eggs, is described as a bet-hedging strategy [8, 11] in plants [12, 18, 30], invertebrates, *e.g*., Daphnia [9], and microorganisms [25] to buffer against environmental variability. Bet-hedging is widely defined as an evolutionary stable strategy in which adults release their offspring into several different environments, here specifically with dormancy at different generations in time, to maximize the chance of survival and reproductive success, thus magnifying the evolutionary effect of good years and dampening the effect of bad years [8, 11]. Dormancy and quiescence sometimes have surprising and counterintuitive consequences, similar to diffusion in activator-inhibitor models [16]. In the following study, we focus more specifically on the evolution of dormancy in plant species [12, 18, 30], but the theoretical models also apply to microorganisms and invertebrate species [9, 25].

Seed banking is a specific life-history characteristic of most plant species, which produce seeds remaining in the soil for short to long periods of time (up to several generations), and it has large but yet underappreciated consequences [11] for the evolution and conservation of many plant species.

First, polymorphism and genetic diversity are increased in a plant population with seed banks compared to the situation without banks. This is mostly due to storage of genetic diversity in the soil [19, 26]. Seed banks also damp off the variation in population sizes over time [26]. Under unfavourable conditions at generation *t*, the small offspring production is compensated at the next generation *t* + 1 by individuals from the bank germinating at a given rate. Under the assumption of large seed banks, the observed population sizes between consecutive generations (*t* and *t* + 1) may then be uncoupled.

Second, seed banks may counteract habitat fragmentation by buffering against the extinction of small and isolated populations, a phenomenon known as the “temporal rescue effect” [7]. Populations which suffer dramatically from events of decrease in population size can be rescued by seeds from the bank. Improving our understanding of the evolutionary conditions for the existence of long-term dormancy and its genetic underpinnings is thus important for the conservation of endangered plant species in habitats under destruction by human activities.

Third, germ banks influence the rate of natural selection in populations. On the one hand, seed banks promote the occurrence of balancing selection for example for color morphs in *Linanthus parryae* [31] or in host-parasite coevolution [27]. On the other hand, the storage effect is expected to decrease the efficiency of positive selection in populations, thus natural selection, positive or negative, would be slowed down by the presence of long-term seed banks. Empirical evidence for this phenomenon has been shown [17], but no quantitative model exists so far. In general terms, understanding how seed banks evolve, affect the speed of adaptive response to environmental changes, and determine the rate of population extinction in many plant species is of importance for conservation genetics under the current period of anthropologically driven climate change.

Two classes of theoretical models have been developed for studying the influence of seed banks on genetic variability. First, Kaj *et al.* [19] have proposed a backward in time coalescent seed bank model which includes the probability of a seed to germinate after a number of years in the soil and a maximum amount of time that seeds can spend in the bank. Seed banks have the property to enhance the size of the coalescent tree of a sample of chromosomes from the above ground population by a quadratic factor of the average time that seeds spend in the bank. This leads to a rescaling of the Kingman coalescent [23] because two lineages can only coalesce in the above-ground population in a given ancestral plant. The consequence of longer seed banks with smaller values of the germination rate is thus to increase the effective size of populations and genetic diversity [19] and to reduce the differentiation among populations connected by migration [32]. This rescaling effect on the coalescence of lineages in a population has also important consequences for the statistical inference of past demographic events [35]. In practice this means that the spatial structure of populations and seed bank effects on demography and selection are difficult to disentangle [5]. Nevertheless, Tellier *et al.* [28] could use this rescaled seed bank coalescent model [19] and Approximate Bayesian Computation to infer the germination rate in two wild tomato species *Solanum chilense* and *S. peruvianum* from polymorphism data [29].

A second class of models assumes a strong seed bank effect, whereby the time seeds can spend in the bank is very long, that is longer than the population coalescent time [14], or the time for two lineages to coalesce can be unbounded. This latest model generates a seed bank coalescent [2], which may not come down from infinity and for which the expected site-frequency spectrum (SFS) may differ significantly from that of the Kingman coalescent [4]. In effect, the model of [19] represents a special case, also called a weak seed bank, where the time for lineages to coalesce is finite because the maximum time that seeds can spend in the bank is bounded.

In the following we mainly have the weak seed bank model in mind where the time in the seed bank is bounded to a small finite number assumed to be realistic for most plant species [12, 18, 29, 30]. Even if we allow for unbounded times a seed may be stored within the soil, we assume that the germination probability decreases rapidly with age such that e.g. the expected time a seed rests in the soil is finite. We develop a forward in time diffusion for seed banks following a Fisher-Wright model with random genetic drift and selection acting on one of two genotypes. The time rescaling induced by the seed bank is shown to be equivalent for the Fisher-Wright and the Moran model. We provide the first theoretical estimates of the effect of seed bank on natural selection by deriving the expected SFS of alleles observed in a sample of chromosomes and the time to fixation of an allele. The main difficulty in the present paper is the non-Markovian character of seedbank models (with the exception of a geometric survival distribution for seeds, in which case the model can be reduced to a Markovian model, see below). The way to deal with this non-Markovian character is based on a separation of time scales. The genetic composition of the population only changes on a slow, so-called evolutionary time scale (thousands of generations), while being fairly stable on a fast, ecological time scale (tens of generations). We assume seeds to have a life span corresponding to this ecological time scale, and thus the seedbank tends to a quasi-stationary state. The non-Markovian character of the model is visible at the ecological time scale, while it vanishes on the evolutionary time-scale due to the quasisteady-state assumption. In other words we ensure the separation of time scales by assuming that most seeds die after a few generations. We demonstrate thereafter that seed banks affect selection and genetic drift differently.

## MODEL DESCRIPTION

We consider a finite plant-population of size *N*. The plants appear in two genotypes *A* and *a*. We assume non-overlapping generations. Let *X*_*n*_ denote the number of type-*A* plants in generation *n* (that is, the number of living type-a plants in this generation is *N* – *X*_*n*_). Plants produce seeds. The number of seeds is assumed to be large, such that noise in the seed bank does not play a role (therefore we call the seed bank “deterministic”). The amount of seeds produced by type-*A*-plants in generation *n* is *β*_*A*_*X*_*n*_, that of type-a plants *β*_*a*_(*N* – *X*_*n*_). The seeds are stored *e.g*. in the soil; some germinate in the next generation, some only in later generations, and some never.

To obtain the next generation of living plants *X*_*n*_, we need to know which seeds are likely to germinate. Let *b*_*A*_(*i*) be the fraction of type-*A* seeds of age *i* able to germinate, and *b*_*a*_(*i*) that of type-a seeds. Hence, the total amount of type-*A* seeds that is able to germinate is given by

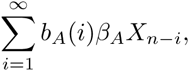

and accordingly, the total amount of all seeds that may germinate

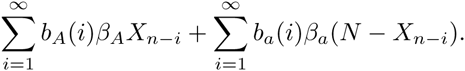

The probability that a plant in generation *n* is of phenotype *A* is given by the fraction of type-*A* seeds that may germinate among all seeds that are able to germinate. The frequency process of the di-allelic Fisher-Wright model with deterministic seed bank reads

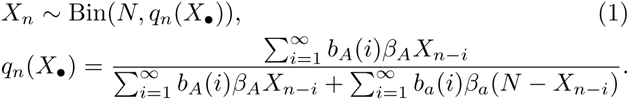

Next we introduce (weak) selection. The fertility of type-a plants is given by

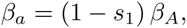

such that *s*_1_ = 0 corresponds to the neutral case. Furthermore, the fraction of surviving seeds is affected. We relate *b*_*a*_(*i*) to *b*_*A*_(*i*) by

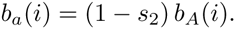

Of course, *s*_2_ has to be small enough to ensure that *b*_*a*_(*i*) ∈ [0,1]. There are other ways to incorporate a fitness difference in the surviving probabilities of seeds, but we feel that this is the most simple version. If we lump *s*_1_ and *s*_2_ in one parameter that scales in an appropriate way for selection,

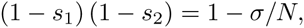

(the sign is chosen in such a way that genotype A has an advantage over genotype a for*σ* > 0 and a disadvantage if *σ* < 0) then eqn. (1) for *q*_*n*_(*X*_•_) with selection becomes

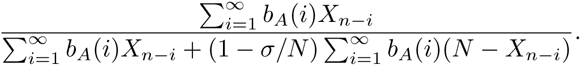

As this ratio is homogeneous of degree zero in *b*_*A*_, we assume 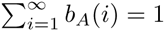. That is, *b*_*A*_(*i*) is considered a probability distribution for the survival of a (type-*A*) seed. We assume that the average life time of a seed is finite,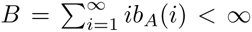. We will implicitly assume that *b*_*A*_(*i*) converge fast enough to zero, such that the separation of ecological and evolutionary time scale is still true. The sum 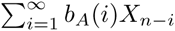 is a moving average. We emphasize this fact by introducing the operator

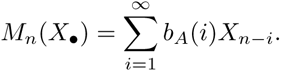

As a consequence, we have *M*_*n*_(*N*) = *N*, and

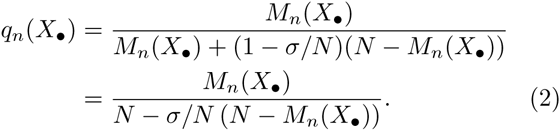

## DIFFUSION LIMIT – GEOMETRIC CASE

As indicated above, if *b*_*A*_(*i*) follow a geometric distribution, then the non-Markovian model introduced above can be reduced to a Markovian model: it is not necessary to track the age of a seed, as all seeds independent of their age have the same mortality resp. germination probability. In this case, and without selection (*σ* = 0), it is straight forward to obtain a diffusion limit that describes the model well on the evolutionary time scale if the population size is (finite but) large. In particular, the diffusion limit is the diffusive Moran model, where we already obtain a first indication how the scaling is affected by the seedbank. Note that the backward process has been analyzed in [3]. This neutral case with a geometric germination rate serves as a warm-up before investigating the full model.

### The Fisher-Wright model without selection

We recall briefly the procedure to derive the diffusion limit for the standard Fisher-Wright model (without seed bank).

- *Model: X*_*n*+1_ ~ Bin(*N*, *X*_*n*_/*N*).
- *Rescale population size:* Let *x*_*n*_ = *X*_*n*_/*N*. Then, *X*_*n*+1_ ~ Bin(*N*, *x*_*n*_). For *N* large, the Binomial distribution approximates a normal distribution with expectation *x*_*n*_ *N* and variance *x*_*n*_(1 – *x*_*n*_)*N*. Let *η*_*n*_ be i.i.d. *N*(0,1)-random variables. Then,

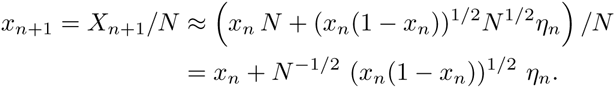
- *Rescale time:* Now define Δ*τ* = 1/*N*, introduce the time *τ* = *n*Δ*τ*, let *u*_*n*Δτ_ = *x*_*n*_, and rescale the index of the normal random variables, that is, replace *η*_*n*_ by *η*_*n*Δ*τ*_ = *η*_*τ*_. Then, *u*_*τ* + Δ*τ*_ − *u*_*τ*_ = Δ*τ*^1/2^ (*u*_*τ*_(1 – *u*_*τ*_))^1/2^ *η*_*τ*_. According to the Euler-Maruyama formula (see *e.g*. [24]), we approximate the diffusive Moran model for *N* large (that is, Δ*τ* = 1/*N* small)

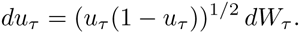

where *W*_*t*_ indicates the Brownian motion.

### Seed bank model with a geometric germination rate and without selection

In the present section we assume that there is no selection (*σ* = 0), and *b*(*i*) follow a geometric distribution with parameter *μ* ∈ (0,1), *b*(1) = *μ* and *b*(*i*) = (1 – *μ*)*b*(*i* – 1). In this case, the delay-model is equivalent to a proper Markov chain.

- *Reformulation of the model:* Define 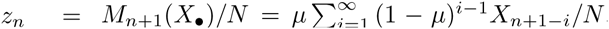. We immediately obtain

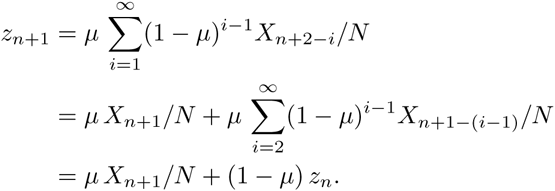 Next (and with the nomenclature of (2)), we have *q*_*n*+1_(*X*_•_) = *M*_*n*+1_(*X*_•_/*N*) = *z*_*n*_. All in all, we reformulated model (1) in the present situation as

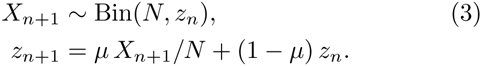 Note that *z*_*n*_ can be interpreted as the state of the seed bank (the fraction of type-*A* seeds that are able to germinate).
- *Rescale population size:* As this model is Markovian, it is simple to derive the diffusion limit. As usual, we start off by defining *x*_*n*_ = *X*_*n*_/*N*, and obtain *z*_*n*_ = *μ x*_*n*_ + (1 – *μ*) *z*_*n* – 1_, *X*_*n*+1_ = Bin(*N*, *z*_*n*_). Approximating the Binomial distribution by a normal distribution for *N* large yields

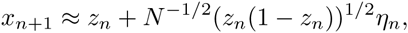 where the *η*_*n*_ ~ *N*(0,1) i.i.d‥ As *x*_*n* + 1_ can be expressed by *z*_*n*_ and *z*_*n* + 1_, the foregoing two equations give

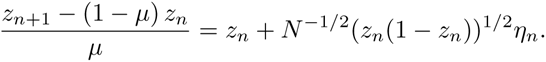 Therefore, *z*_*n*+1_ – *z*_*n*_ = *μN*^−1/2^ (*z*_*n*_(1 – *z*_*n*_))^1/2^ *η*_*n*_.
- *Rescale time:* Scaling time by *N* yields for *u*_*n*_/*N* = *z*_*n*_ and *τ* = *n*/*N*

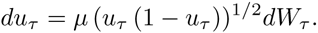 If we define *B* = 1/*μ* (the expected value of a geometric distribution with parameter *μ*), we may write this equation as

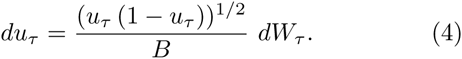 We find a diffusive Moran model for the state of the seed bank with rescaled time scale. The factor 1/*B* has been already proposed in the paper of Kaj, Krone and Lascoux [19], who analyzed a seedbank process backward in time.

## DIFFUSION LIMIT – GENERAL CASE

We expect a similar result as above to hold in the general case. A difference between the two cases is that we naturally considered the state of the seed bank before, while in the general case we will focus on the state of living plants. As discussed before, the center of the analysis below is an additional step that investigates the quasi-stationary state of the seedbank at evolutionary time scale; this additional step is necessary to deal with the non-Markovian character of our model.

### Rescale population size

From (2), we immediately have

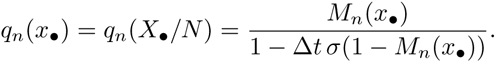

Using Normal approximation of the Binomial distribution leads to

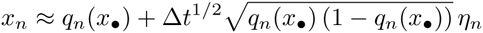

where *η*_*n*_ ~ *N*(0,1) are i.i.d‥ Taylor expansion of *q*_*n*_(*x*_•_) w.r.t. Δ*t* yields in lowest order

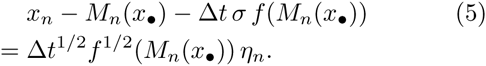

with *f*(*x*) = *x*(1 – *x*).

### Perturbation approach

The leading term of eqn. (5) is *x*_*n*_ – *M*_*n*_(*x*_•_). This difference must not become too large, as all other terms in the equation are at least of order Δ*t*^1/2^. That is, the state *x*_*n*_ can only slowly drift away from *M*_*n*_(*X*_•_) (which represents the state of the seed bank). Hence, for a reasonable number of time steps (on the ecological time scale), *M*_*n*_(*X*_•_) is fairly constant. In order to disentangle the evolutionary and the ecological time scale, we introduce *ε* = Δ*t*^1/2^, expand *x*_*n*_ w.r.t. *ε*,

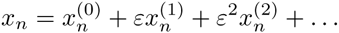

and rewrite eqn. (5) as

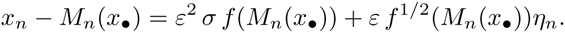

Taylor expansion and equating equal powers of *ε* yields

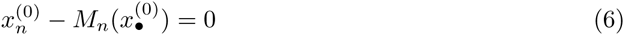

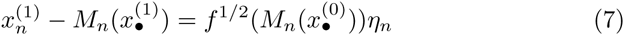

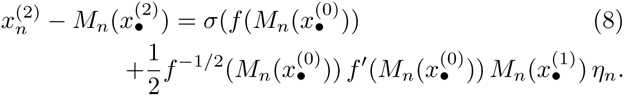

**Zero order:** The zero order term 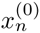 follows a deterministic dynamics. As *M*_*n*_ is an averaging operator the solution becomes constant in the long run. The system saddles on the slow manifold, consisting of constant sequences. At this point it is important that *b*_*A*_(*i*) tend fast enough to zero, s.t. 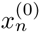 indeed approximates on the fast (ecological) time scale the slow manifold. We assume 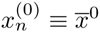.

**First order:** The recursive equation (7) is well known as an auto-regression (AR) model in the statistical modeling of time series [6]. We define 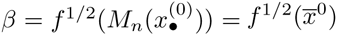(note that *β* is a real number and not a random variable) and convert the AR model into a moving average equation. Thereto we introduce the back-shift operator acting on the index of a sequence, *Lz*_*n*_ = *z*_*n*–1_, and the power series

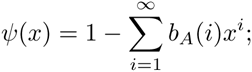

Eqn. (7) becomes in this notation

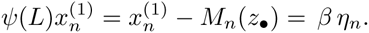

Note that *ψ*(1) = 0, which does mean that the AR model is non-stationary (this process is also called an ARIMA model for time series [6, Chapter 9]). We do not find a power series *ψ*^*^(*x*) well defined at *x* = 1 such that *ψ*^*^(*x*) *ψ*(*x*) = 1. Therefore, we rewrite *ψ*(*x*) as *ψ*(*x*) = (1 – *x*) *ψ*̃(*x*) (which is the defining equation of *ψ*̃(*x*)). As

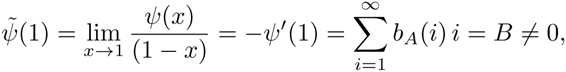

we do find *ψ*^*^(*x*) such that *ψ*^*^(*x*)*ψ*̃(*x*) = 1, and hence *ψ*^*^(*x*)*ψ*(*x*) = 1 – *x* in a neighbourhood of *x* =1. As an immediate consequence (used later) we have *ψ*^*^(1) = 1/*B*. If we multiply the equation 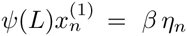, we obtain

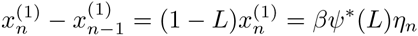

and

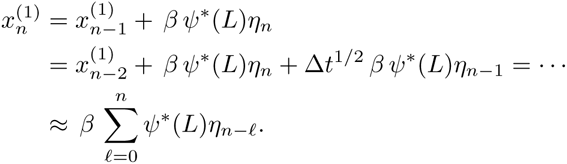

Let 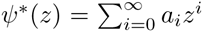. We expand the sum above, and obtain

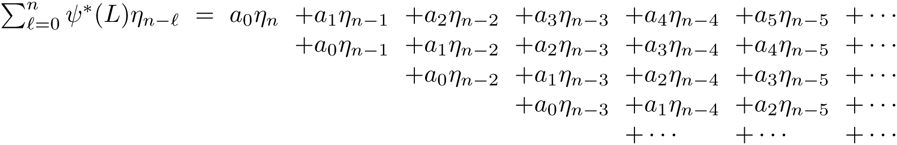

If we inspect not rows (that have *ψ*^*^(*L*)*η*_*i* – ℓ_ as entries) but columns (that contain always the same random variable *η*_*i* – ℓ_), we find that the coefficient in front of one given random variable *η*_*i* – ℓ_ approximates *ψ*^*^(1) for *ℓ* → ℞.

At this point, we want to write 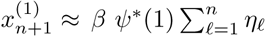.

This is only true, also in an approximate sense, if *n* is large and the state *x*_*n*_ does hardly change over a time scale that allows 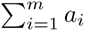 to converge to *ψ*^*^(1) = 1/*B*. If Δ*t*^1/2^ is small, then indeed *x*_*n*_ ≈ *x*̄_0_ on the ecological time scale, as required. Hence, for Δ*t* small we are allowed to assume

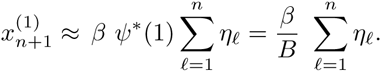

Thus 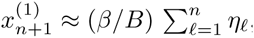, and for *n* large

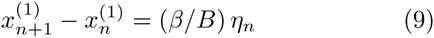

where, as before, *η*_*n*_ ~ *N*(0,1) i.i.d‥

**Second order:** With 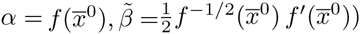, we may write

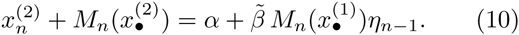

If 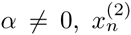 incorporates a deterministic trend. We first remove this trend defining 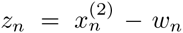 with *ω*_*n*_ = *nα*/*B*. Then, 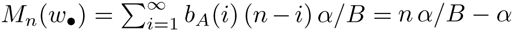, and, with 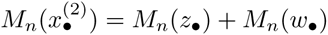,

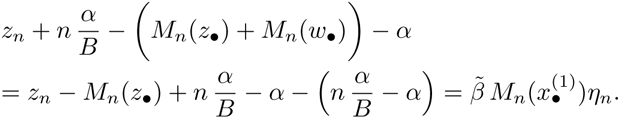

We obtain an AR model for *z*_*n*_ without trend,

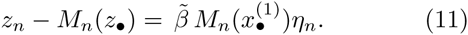

It turns out, that we need not to analyze *z*_*n*_ in detail. It is sufficient to note that *z*_*n*_ is a random variable with expectation zero.

**Result:** All in all, we conclude

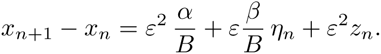

We only take into account the lowest order in the deterministic drift resp. in the random perturbations. As *ε* (*β*/*B*) *η*_*n*_ dominates *ε*^2^ *z*_*n*_, we drop the latter term, replace in *α*, *β* the variable *x*̄^0^ by *x*_*n*_, and end up with

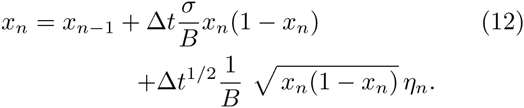

**Numerical simulation:** We compare the result of these computations with numerical simulations. Thereto we consider the linear model

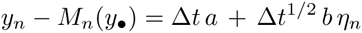

with *a*, *b* ∈ ℝ. If *y*_*n*_ = 0 for *n* ≤ 0, we expect that *y*_*n*_ (for *n* ≥ 1) approximately to satisfy

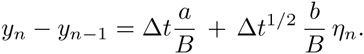

That is, *y*_*n*_ is approximately normally distributed with expectation *n*Δ*ta*/*B*, and variance *n*Δ*tb*^2^/*B*^2^. For simulations, we choose *a* = 1, Δ*t* = 0.01, *b* = 2 and 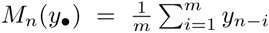 for *m* = 9, that is, *B* = 5.

The simulations show an excellent agreement with our computations (Figure 1).

**FIG. 1.**
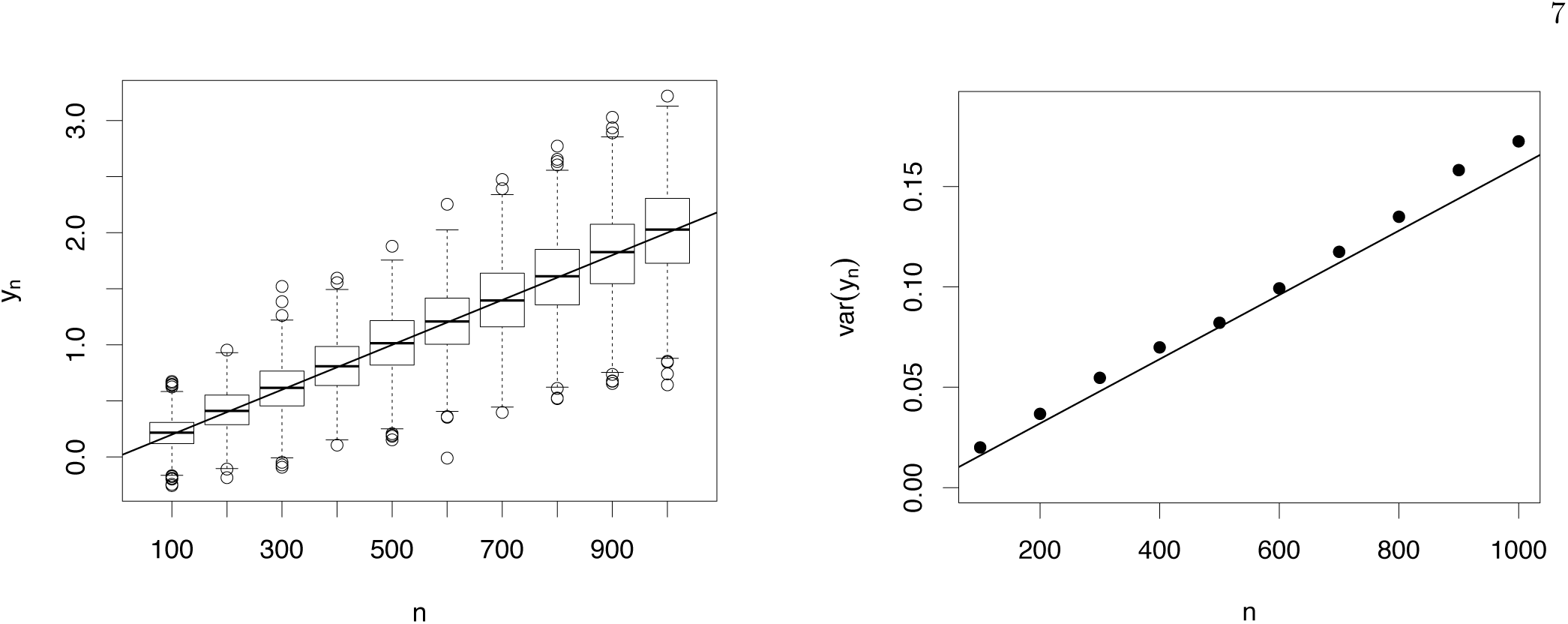
Simulation of the AR model (1000 runs). Samples have been taken at time steps 100, 200,…, 1000. (left) Boxplot of the simulated time series *y*_*n*_ at indicated time points together with the mean according to eqn. 10 (line). (right) Variance of the simulated time series at indicated time points (dots), together with the variance according to eqn. 10 (line). For parameters used: see text.

### Rescale time

As before, we define *u*_*n*Δ*t*_ = *x*_*n*_, and use the Euler-Maruyama-formula to conclude that *u*_*t*_ approximates for Δ*t* → 0 the stochastic differential equation

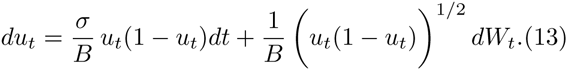

Please note that this result seems to inherit the usual stability of a diffusion limit w.r.t. the detailed model assumptions: if we start off with a Moran model instead of a Fisher-Wright model combined with a seed bank, we again obtain a diffusion limit of similar form (see Appendix).

We now change the time scale such that the variance coincides with the standard diffusive Moran model. If we define *τ* = *t*/*B*^2^, then the SDE reads

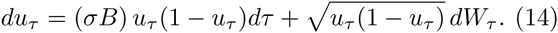

**Scaling of the selection parameter.** We conclude, in line with previous findings (see discussion), that the appropriate scaling of time for the Fisher-Wright model with seed bank is not 1/*N* but 1/(*B*^2^ *N*). Moreover, the effective selection rate (w.r.t. this time) is increased by the average number of generations *B* the seeds sleep in the soil.

## THE FORWARD DIFFUSION EQUATION FOR SEED BANK MODELS WITH SELECTION

In analogy to above, we consider a single locus and two allelic types *A* and a with frequencies *x* and 1 – *x*, respectively, at time zero. Time is scaled in units of 2*N* generations. In the diffusion limit, as *N* → ∞, the probability *f*(*y*, *t*)*dy* that the type-*A* genotype has a frequency in (*y*, *y* + *dy*) is characterized by the following forward equation (see [20] for *B* = 1):

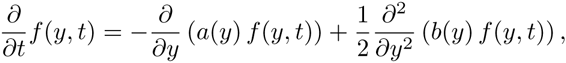

where the drift and the diffusion terms are given by *a*(*y*) = *σ y*(1 – *y*)/*B* and *b*(*y*) = *y*(1 – *y*)/*B*^2^, respectively.

For the derivations of the frequency spectrum and the times to fixation we require the following definitions. The scale density of the diffusion process is given by

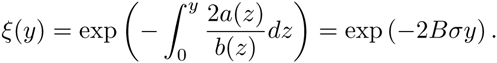

The speed density is obtained (up to a constant) as

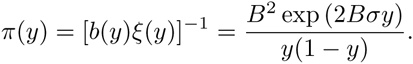

The probability of absorption at *y* = 0 is given by

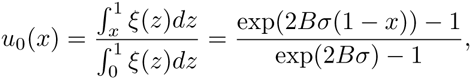

and *u*_1_(*x*) = 1 – *u*_0_(*x*) gives the probability of absorption at *y* = 1.

### Site-frequency spectra

The site-frequency spectrum (SFS) of a sample (*e.g*., [10, 15, 34]) is widely used for population genetics data analysis. A sample of size k is sequenced, and for each polymorphic site the number of individuals in which the mutation appears is determined. In this way, a dataset is generated that summarizes the number of mutations *ζ*_*k*, *i*_ appearing in *i* individuals, *i* = 1,…, *k* – 1. That is, *ζ*_*k*, 1_ = 10 indicates that 10 mutations only appeared once, and *ζ*_*k*, 2_ = 5 tells us that five mutations were present in two individuals (where the pair of individuals may be different for each of the five mutations). Note that neither *ζ*_*k*, 0_ nor *ζ*_*k*, *k*_ are sensible: a mutation that appears in none or all individuals of the sample cannot be recognized as a mutation. In practice, it is often not possible to know the ancestral state. Then the folded SFS *η*_*k*, *i*_ = (*ζ*_*k*, *i*_ + *ζ*_*k*, *k* – *i*_)(1 + 1_{*i*=*k*–*i*}_)^−1^ can be used. Since both empirical observations and theoretical results for the folded SFS follow instantaneously from the unfolded one, we only consider the unfolded version.

For the derivation of the theoretical SFS, we assume that mutations occur according to the infinitely-many sites model [21]. The scaled mutation rate is given by *θ* = 4*N v*, where *v* is the mutation rate per generation at independent sites. Assuming that each mutant allele marginally follows the diffusion model specified above, the proportion of sites where the mutant frequency is in (*y*, *y* + *dy*) is given by [15]

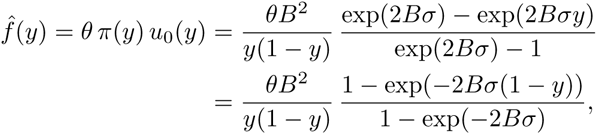

where *f*̂(*y*) denotes the equilibrium solution of the population SFS. For neutrality, we immediately obtain *f*̂(*y*) = *θ B*^2^/*y* by letting a *σ* 0 in the foregoing equation.

The equilibrium solution of the SFS for a sample of size *k* is obtained via binomial sampling (see [33] for *B* = 1) as

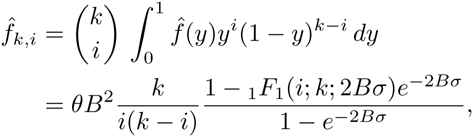

where _1_*F*_1_ denotes the confluent hypergeometric function of the first kind [1]. For neutrality, we again immediately obtain *f*̂_*k*, *i*_ = *θ B*^2^/*i* by letting *σ* → 0. For a large number of mutant sites, the relative SFS 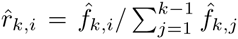 approximates the empirical distribution 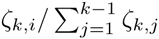 for a constant population size. Note that the solutions for the absolute SFS assume that mutations can occur at any time. When assuming that mutations can only arise in living plants [19], *θ* has to be replaced by *θ*/*B* in the respective equations. Both mutation models give equivalent results for the relative SFS.

As shown in Figure 2 (left), the neutral diffusion approximation is in line with the simulation results of the original discrete model. The theoretical relative SFS for a sample of 250 individuals approximates the simulated SFS, which is obtained as an average over 10,093 repetitions. In every iteration, the sample is drawn from an initially monomorphic population of 1000 individuals after 400,000 generations (so that the population has reached an equilibrium). Figure 2 (right) illustrates the enhanced effect of selection proportional to the length of the seed bank.

**FIG. 2.**
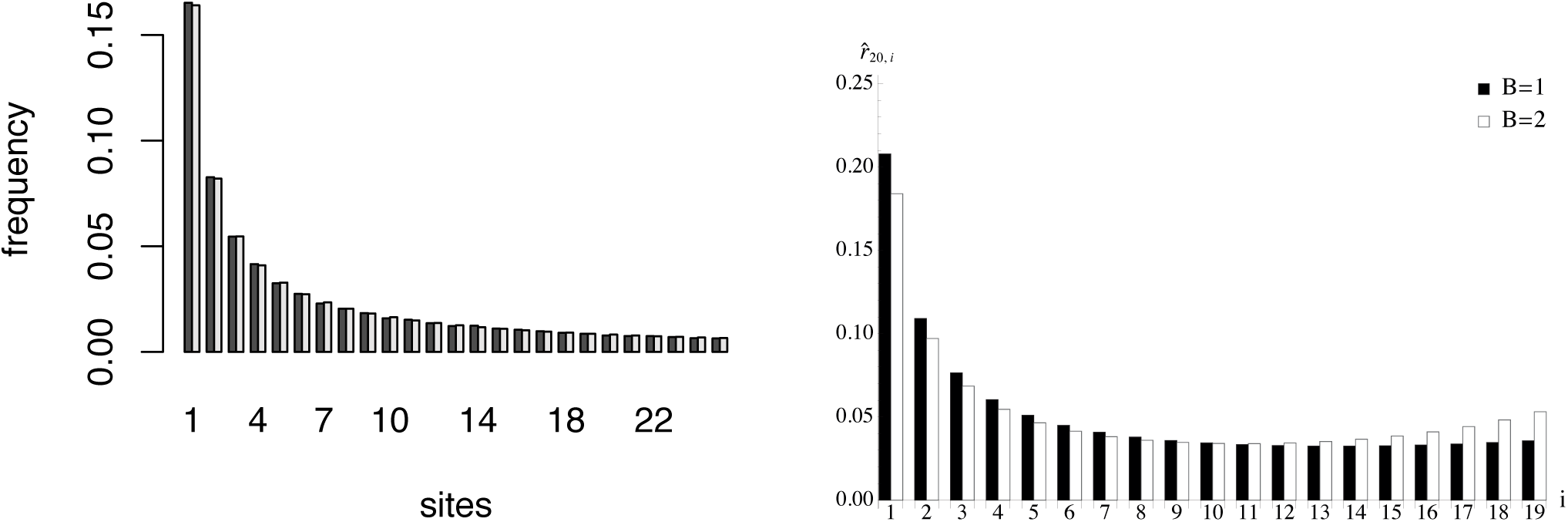
(left) Simulation and theoretical prediction for the neutral relative SFS and a uniformly distributed seed bank of length *B* = 10. For the simulation of the original discrete model the population size was chosen as 1000, we started without mutations and stopped the process after 400,000 generations to calculate the SFS as an average over 10,093 repetitions. The light gray bar shows the theoretical result, the dark gray bar shows the simulation outcome. In both cases a sample of 250 individuals was drawn. (right) Theoretical results for the relative SFS of a sample of size 20 are plotted for positive selection of strength *σ* = 2 without (*B* = 1) and with a seed bank of length *B* = 2.

### Times to fixation

We assume that both *y* = 0 and *y* = 1 are absorbing states and start by considering the mean time until one of these states is reached in the diffusion process specified above. The mean absorption time *t*̄ can be expressed as [13]

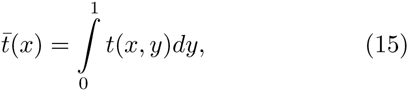

where

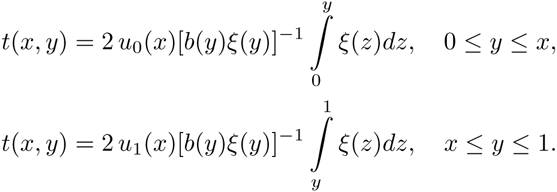

For genetic selection the integral in (15) cannot be analytically solved. For selective neutrality, we obtain *t*̄(*x*) = –2*B*^2^ (*x* log(*x*) + (1 – *x*) log(1 – *x*)) (see *e.g*. [13] for *B* = 1) by employing the drift term, the scale density and the probabilities of absorption as specified above. Now, we evaluate the time until a mutant allele is fixed conditional on fixation as 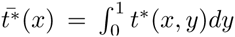, where *t*^*^(*x*, *y*) = *t*(*x*, *y*) *u*_1_(*y*)/*u*_1_(*x*). For genic selection the mean time to fixation in dependency of x can only be derived as a very lengthy expression in terms of exponential integral functions. The neutral result is found as *t*̄(*x*) = –2*B*^2^(1 – *x*)/*x* log(1 – *x*) and in accordance with a classical result [22] for *B* = 1. For *x* → 0, we obtain

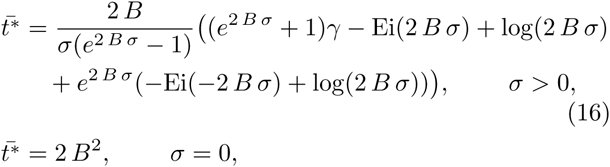

where *γ* is Euler’s constant and Ei denotes the exponential integral function [1].

In Figure 3 (left), we compare the time to absorption of the original discrete seed bank model by means of simulations with the theoretical result obtained from the diffusion approximation. For *b*_*A*_ we use uniform distributions, where we vary the expected values between 1 and 8 corresponding to the length of the seed banks between 1 and 15. We choose an initial fraction of 0.5 for the type-*A* genotypes. The simulations show a good agreement between our analytical approximation and the numerical simulations. In Figure 3 (right), we show the effect of the seed bank on the times to fixation conditional on fixation of the type-A genotype for neutrality and positive selection.

**FIG. 3.**
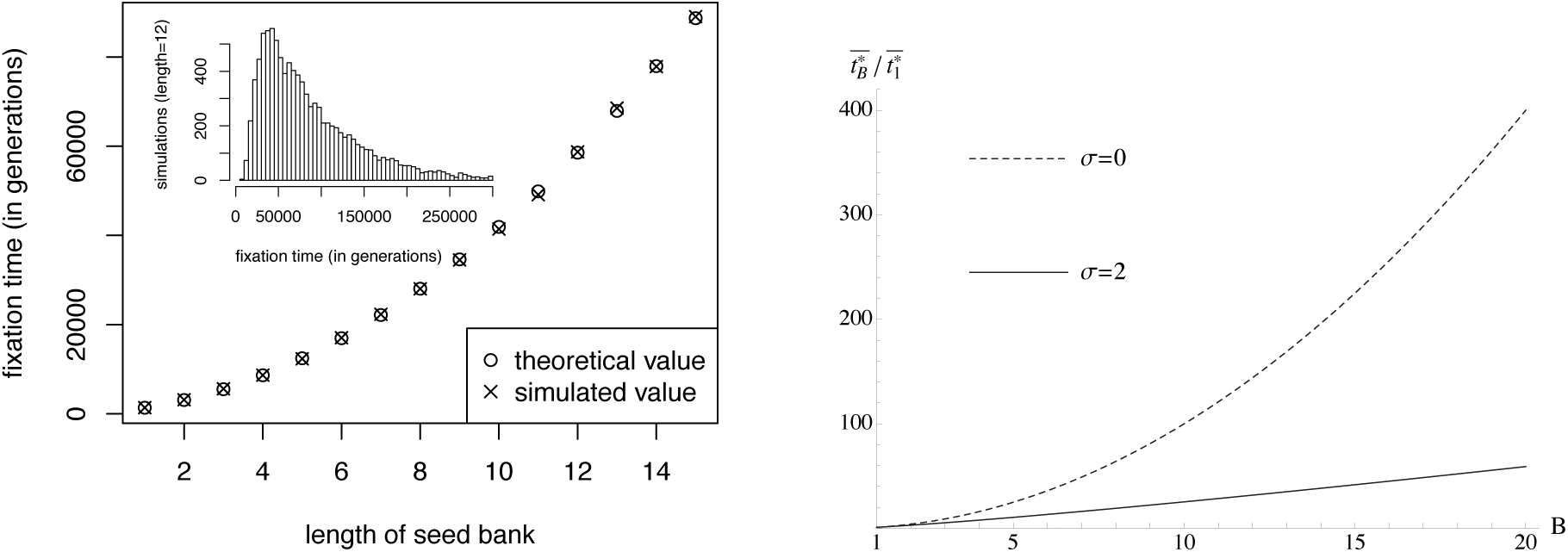
(left) Simulation and theoretical prediction for the time to fixation of a seed bank model. The population size is 1000 and 50% of the individuals are initially of genotype A. We simulated 10,000 runs for each mean value. The simulated distribution of the time to fixation is shown in the histogram at the upper left corner taking the data of the simulated seed bank of length *B* = 12. (right) The ratios of the conditional fixation times with and without seedbank are plotted against the length of the seed bank B for neutrality and selection by employing (16). The additional index in the ratio is used to formally distinguish the cases with and without seed bank.

## DISCUSSION

Within this study, we develop a forward in time Fisher-Wright model of a deterministically large seed bank with drift occurring in the above-ground population. The time that seeds can spend in the bank is bounded and finite, as assumed to be realistic for many plant or invertebrate species. We demonstrate that scaling time in the diffusion process by a factor *B*^2^ generates the usual Fisher-Wright time scale of genetic drift with *B* being defined as the average amount of time that seeds spend in the bank. The conditional time to fixation of a neutral allele is slowed down by a factor *B*^2^ (Figure 3 (right), dotted line) compared to the absence of seed bank. These results are consistent with the backward in time coalescent model from Kaj *et al.* [19], and differs from the strong seed bank model of Blath *et al.* [2]. We evaluate the SFS based on our diffusion process and confirm agreement to the SFS obtained under discrete time Fisher-Wright simulations.

In the second part of the study, we introduce selection occurring at one of the two alleles, mimicking positive or negative selection. Two features of selection under seed banks are noticeable. First, selection is slower under longer seed banks (Figure 3 (right), solid line) confirming previous intuitive expectations [17]. Second, when computing the SFS with *B* = 2 and without seed bank (*B* = 1) under positive selection (*σ* = 2) we reveal a stronger signal of selection for the seed bank by means of an amplified uptick of high-frequency derived variants. This effect becomes more prominent with longer seed banks and also holds for purifying selection, under which an increase in low-frequency derived variants is induced by the seed bank. We explain this counterintuitive results as follows: longer seed banks increase, on the one hand, the selection coefficient a generating *σ* stronger signal at equilibrium (Figure 2 (right)), and on the other hand, the time to reach this equilibrium state (Figure 3 (right)). Our predictions are consistent with the inferred strengths of purifying selection in wild tomato species. Indeed, purifying selection at coding regions appears to be stronger in *S. peruvianum* than in its sister species *S. chilense* [28] with *S. peruvianum* exhibiting a longer seed bank [29].

*This research is supported in part by Deutsche Forschungsgemeinschaft grants TE 809/1 (AT) and STE 325/14 from the Priority Program 1590 (DZ)*.

## Appendix: Moran model with deterministic seed bank

We briefly sketch the arguments that allow to handle a Moran model with seed bank; the reasoning is completely parallel to the time-discrete case. In order to keep this appendix short, we do not take into account selection but focus on the neutral model.

### Model

We start off with the individual based model. Let the population size be *N*, *X*_*t*_ the number of genotype-A-plants, *δ* the death rate, and *b*(*s*) the distribution of the ability for a seed at age *s* to germinate; we require 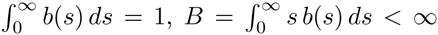, and *b*(*s*) sufficiently smooth. Then, the rate for the transition *X*_*t*_ → *X*_*t*_ + 1 is given by

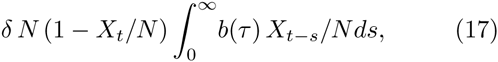

while that for a decrease of X_t_ by 1 reads

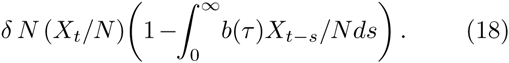

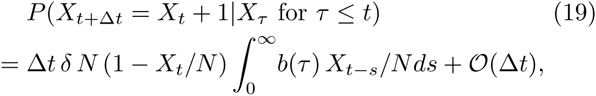

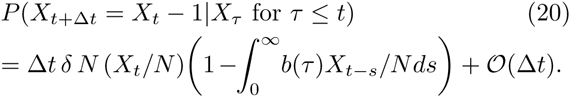

Note that the delay process requires the knowledge of the complete history {*X*_*s*_}_*s* < *t*_. The usual continuous limit for *x*_*t*_ = *X*_*t*_/*N* yields (with *ε* = 1/*N*)

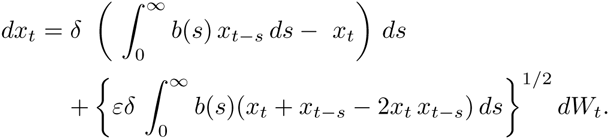

If we rescale time in the usual way, *τ* = *εt*, and define 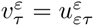, we obtain

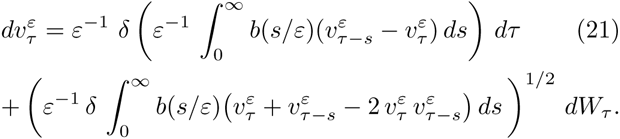

The aim here is to find heuristic arguments indicating that 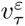 approximates for *ε* → 0 the solution of a Moran diffusion process with rescaled time, paralleling equation (13).

Note that, in some sense, the terms in this time-continuous model are better to interpret than the parallel terms in the Fisher-Wright model: both terms within the brackets are moving averages, and clearly

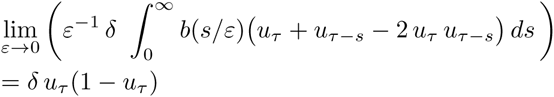

for a function *u*_*τ*_ that is reasonably smooth. For the drift term, we find similarly

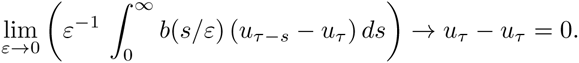

However, in eqn. (21), this bracket is divided by *ε*, and hence does not vanish for *ε* → 0. If we take a closer look, we find that a deviation of *x*_*τ*_ from the moving average (the state of the seed bank) is punished. That is, the state of living plants can change only slower in comparison with a model without seed bank, and therefore for *ε* → 0 we expect a diffusion model at a slower time scale.

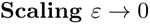

We drop the superscript *ε* in 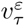, and write simply *υ*_*τ*_. In order to use the arguments developed above, we discetize the stochastic differential-delay equation by the Euler-Maruyama formula, and find

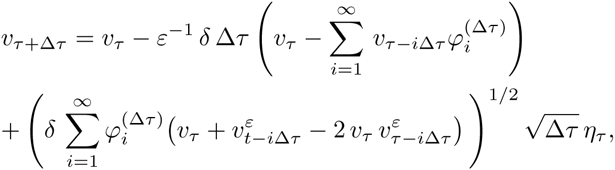

where *η*_*τ*_ are i.i.d. *N*(0,1) distributed, and the weights 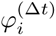 are chosen as

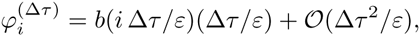

such that 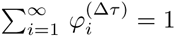. If we now define

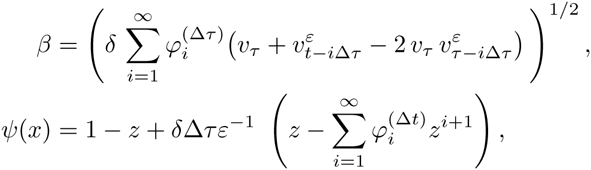

we may rewrite the discretized equation for *υ*_*τ*_ as

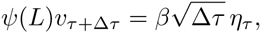

where *Lυ*_*τ*_ = *υ*_*τ* – Δ*τ*_. We are now in the position to apply the computations about the quasi-stationary state of the seedbank (neglecting the time-dependency of *β*). As

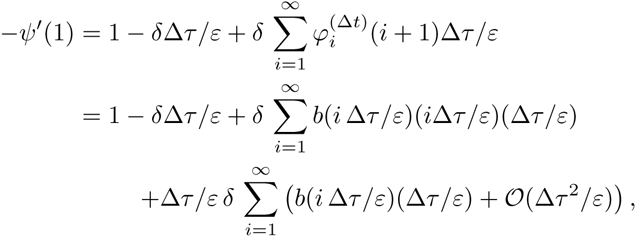

we have

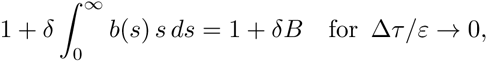

and conclude that approximately

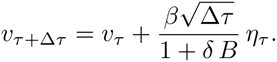

Hence, for *ε* → 0 we expect (according to these heuristic arguments) that 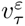 satisfies the rescale diffusion equation

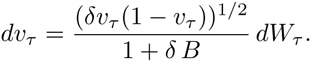

If we define *G* = 1/*δ*, the average inter-generation time of living plants, this equation becomes even closer to that derived for the Fisher-Wright case,

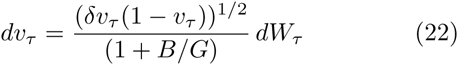

as it becomes clear that the correction factor 1 + *B*/*G* measures the average time a seed rests in the soil in terms of generations.

## References

[1] Abramowitz, M., and I. A. Stegun (1964), Handbook of mathematical functions: with formulas, graphs, and mathematical tables (Dover).

[2] Blath, J., B. Eldon, A. González-Casanova, N. Kurt, and M. Wilke-Berenguer (2015), Genetics 200, 921.

[3] Blath, J., A. González Casanova, N. Kurt, and D. Spanò (2013), J. Appl. Probab. 50, 741.

[4] Blath, J., A. González-Casanova, N. Kurt, and M. Wilke-Berenguer (2016), Ann. Appl. Prob. 26, 857.

[5] Böndel, K. B., H. Lainer, T. Nosenko, M. Mboup, A. Tellier, and W. Stephan (2015), Mol. Biol. Evol. 32, 2932.

[6] Brockwell, P. J., and R. A. Davis (2009), Time Series: Theory and Methods (Springer).

[7] Brown, J. H., and A. Kodric-Brown (1977), Ecology 58, 445.

[8] Cohen, D. (1966), J. Theor. Biol. 12, 119.

[9] Decaestecker, E., S. Gaba, J. A. M. Raeymaekers, R. Stoks, L. Van Kerckhoven, D. Ebert, and L. De Meester (2007), Nature 450, 870.

[10] Etheridge, A. (2011), Some Mathematical Models from Population Genetics, LNM 2012 (Springer).

[11] Evans, M. E. K., and J. J. Dennehy (2005), Q. Rev. Biol 80, 431.

[12] Evans, M. E. K., R. Ferriere, M. J. Kane, and D. L. Venable (2007), Am. Nat. 169, 184.

[13] Ewens, W. J. (2004), Mathematical Population Genetics: I. Theoretical Introduction (Springer).

[14] González-Casanova, A.., E. A. von Wobeser, G. Espín, L. Servín-González, N. Kurt, D. Spanó, J. Blath, and G. Soberón-Chávez (2014), J. Theor. Biol. 356, 62.

[15] Griffiths, R. C. (2003), Theor. Popul. Biol. 64, 241.

[16] Hadeler, K. (2013), J. Math. Biol. 66, 649.

[17] Hairston, N. G., and B. T. Destasio (1988), Nature 336, 239.

[18] Honnay, O., B. Bossuyt, H. Jacquemyn, A. Shimono, and K. Uchiyama (2008), Oikos 117, 1.

[19] Kaj, I., S. M. Krone, and M. Lascoux (2001), J. Appl. Probab. 38, 285.

[20] Kimura, M. (1955), in Cold Spring Harbor Symposia on Quantitative Biology, Vol. 20 (Cold Spring Harbor Laboratory Press) pp. 33–53.

[21] Kimura, M. (1969), Genetics 61, 893.

[22] Kimura, M., and T. Ohta (1969), Genetics 61, 763.

[23] Kingman, J. F. C. (1982), J. Appl. Probab. 19A, 27.

[24] Kloeden, P. E., and E. Platen (1992), Numerical Solution of Stochastic Differential Equations, Applications of Mathematics, Stochastic Modelling and Applied Probability, Vol. 23 (Springer).

[25] Lennon, J. T., and S. E. Jones (2011), Nat. Rev. Microb. 9, 119.

[26] Nunney, L. (2002), Am. Nat. 160, 195.

[27] Tellier, A., and J. K. M. Brown (2009), Am. Nat. 174, 769.

[28] Tellier, A., I. Fischer, C. Merino, H. Xia, L. Camus-Kulandaivelu, T. Stadler, and W. Stephan (2011), Heredity 107, 189.

[29] Tellier, A., S. J. Y. Laurent, H. Lainer, P. Pavlidis, and W. Stephan (2011), Proc. Natl. Acad. Sci. U.S.A. 108, 17052.

[30] Tielbörger, K., M. Petruů, and C. Lampei (2012), Oikos 121, 1860.

[31] Turelli, M., D. W. Schemske, and P. Bierzychudek (2001), Evolution 55, 1283.

[32] Vitalis, R., S. Glemin, and I. Olivieri (2004), Am. Nat. 163, 295.

[33] Živković, D., M. Steinrücken, Y. S. S. Song, and W. Stephan (2015), Genetics 200, 601.

[34] Živković, D., and W. Stephan (2011), Theor. Popul. Biol. 79, 184.

[35] Živković, D., and A. Tellier (2012), Mol. Ecol. 21, 5434.

